# Prenatal cannabinoid exposure affects central cardiorespiratory control in young male and female rats

**DOI:** 10.1101/2025.01.16.633429

**Authors:** Luis Gustavo A. Patrone, Marlusa Karlen-Amarante, Luciane H. Gargaglioni, Daniel B. Zoccal

**Author notes:** Corresponding author: Department of Physiology and Pathology, School of Dentistry of Araraquara, São Paulo State University (UNESP), Rua Humaitá, 1680, 14801-903, Araraquara, São Paulo, Brazil. Dalton Cardiovascular Research Center, University of Missouri, Columbia, MO, USA. joint senior authors (Luciane H. Gargaglioni, Daniel B. Zoccal).

## Abstract

Cannabis use among pregnant women is rising globally, mainly for recreational and medical reasons to relieve symptoms like nausea, vomiting, anxiety, and insomnia. This trend is reinforced by the misconception that its natural origin guarantees safety, along with government policies promoting legalization. However, exposure to cannabinoids *in utero* can impact normal offspring’s neurodevelopment and induce malfunctioning of various physiological systems, including the cardiorespiratory function. The present study investigated whether prenatal cannabinoid exposure disrupts the generation and control of autonomic and respiratory activities in early adulthood. Using *in situ* preparations of juvenile male and female rats (27-28 days old) exposed to a synthetic cannabinoid (WIN 55,212-2; 0.5 mg/kg/day, n=4-9) or vehicle (n=3-10) during gestation, we analyzed the activity of nerves innervating respiratory muscles and blood vessels. We noticed that females receiving WIN prenatally exhibited a reduced excitatory drive (post-inspiratory activity, post-I) to laryngeal muscles under resting conditions, suggesting impaired control of upper airway patency. Moreover, males and females exposed to WIN displayed reduced post-I and abdominal expiratory motor activities during stimulation of carotid body chemoreceptors (mimicking low oxygen situations) or exposure to high carbon dioxide levels, indicating an inability to mount appropriate reflex respiratory motor responses during blood gas disturbances. In addition, WIN-treated males showed attenuated sympathoexcitatory responses to carotid body activation or hypercapnia, evidencing a limited capacity to promote sympathetic-mediated hemodynamic changes. Thus, manipulating the fetal endocannabinoid system impacts the development of networks controlling respiratory and autonomic functions, leading to negative, long-term consequences for ventilation and cardiovascular function.

## 1. INTRODUCTION

The development of the central nervous system (CNS) is a highly dynamic process in which the formation of neural components, including the structural maturation and differentiation into specific phenotypes, and their connections emerge from complex interactions among genes, transcriptional regulators, and neurotrophic factors [1]. The endocannabinoid signaling during prenatal development plays a crucial role in the ontogeny of the CNS, regulating neurogenesis, cell differentiation, synaptogenesis, and synaptic plasticity [2–4]. It is also considered an important neuromodulatory pathway driving the development of different neurotransmitters [5–8].

Exposure to adverse conditions *in utero* can reshape ontogenetic development, resulting in epigenetic-dependent deviations from the expected formation and maturation of the CNS and the occurrence of dysfunctions in physiological systems in the offspring. [9–11]. Not exclusively central pathways but peripheral sensory mechanisms are also susceptible to adaptive and maladaptive changes during prenatal development as a result of intrauterine disturbances [12–14]. Experimental evidence indicates that overstimulation of the endocannabinoid system during intrauterine life can be detrimental to physiological functions in postnatal life [15, 16]. These observations are relevant in the context of the rising number of pregnant women consuming cannabis-derived products to manage pregnancy-related conditions (e.g., nausea, weight gain, sleep difficulty) or pre-existing maternal health conditions (e.g., anxiety, depression, chronic pain) [17, 18], aligned with global flexibilization policies of cannabis consumption or cannabinoid-based products use. The lack of deep understanding of the effects of intrauterine cannabinoid exposure on offspring’s physiological functions also limits awareness within the population.

The development of cardiorespiratory control mechanisms *in utero* is subject to exhibiting long-term modifications as a result of exposure to endocannabinoids [19–24]. Experimental studies show that offspring prenatally exposed to the synthetic cannabinoid (WIN 55,212-2) exhibited, during the first weeks of life, an increased number of catecholaminergic neuronal populations and elevated cannabinoid receptor type 1 (CB_1_) expression in the brainstem combined with amplified ventilatory responses during conditions of elevated inspired carbon dioxide (or hypercapnia) [23]. This plasticity in the respiratory homeostatic mechanisms induced by prenatal exposure to WIN persisted until adulthood in a sex-dependent manner, with males showing augmented ventilatory responses to hypercapnia or reduced oxygen levels (hypoxia) and females exhibiting attenuated responses [24]. Additionally, adult animals prenatally exposed to WIN were prone to show elevated arterial pressure and heart rate levels during resting conditions or when subjected to hypoxia, mainly the females [24].

Despite evidence showing alterations in the functioning of the cardiorespiratory system induced by prenatal exposure to cannabinoids, it remains unclear whether these changes are driven by central or peripheral mechanisms. In this context, it is of critical importance to understand the impact of prenatal cannabinoid overstimulation on the motor activity governing pumping and upper airway respiratory muscles as well as on sympathetic activity supplying blood vessels. To address this, the current study investigated the effects of prenatal cannabinoid exposure on the activity of phrenic, abdominal, central vagus, and thoracic sympathetic activities of juvenile/early adult rats. By treating the animals with WIN prenatally, we tested the hypothesis that cannabinoid exposure *in utero* disrupts the functioning of brainstem circuitries controlling cardiorespiratory motor outputs.

## 2. MATERIAL AND METHODS

### 2.1 Animals

Male and female adult Wistar rats (250 – 300 g) were obtained from the Animal Care facility of UNESP – Botucatu, São Paulo, Brazil, and used for mating. The gestational day 0 was defined as when spermatozoa were detected through vaginal wash. Pregnant female rats were used for prenatal treatment, and their juvenile offspring of both sexes (first generation) at 27-28 days old were utilized to obtain *in situ* working heart-brainstem preparations (WHBP). Pregnant rats and their litters were individually housed in cages in a temperature-controlled room maintained at 25 ± 1°C, with a 12:12-h light– dark cycle (lights on at 6:30 a.m.), with access to water and food *ad libitum*. The number of newborns per litter at P0 was set at between 8 and 10 to prevent variations in litter sizes and offspring nutrition during the postnatal period. All pups were born in the Animal Care facility of UNESP – Jaboticabal, São Paulo, Brazil, and stayed in the same cage with their mother until weaning at P21. Afterward, they were grouped based on sex and treatment until they reached the experimental age (P27-28). The juveniles were randomly selected from different litters to mitigate potential litter-related effects. All experiments were performed between 7:00 a.m. and 6:00 p.m. at the Department of Physiology and Pathology of the School of Dentistry at UNESP - Campus Araraquara, São Paulo. The experiments comply with ARRIVE and the National Council of Control in Animal Experimentation (CONCEA-MCTI-Brazil) guidelines and were approved by the Animal Care and Use Committee of the College of Agricultural and Veterinary Sciences (Protocol: 011284/17) and School of Dentistry of Araraquara (Protocol: 31/2016).

### 2.2 Prenatal cannabinoid treatment

Pregnant rats were treated from gestational day 0 to the 21^st^ with either vehicle (dimethyl sulfoxide - DMSO 50%, diluted in sterile water) or synthetic cannabinoid WIN 55,212-2 (Sigma Aldrich, USA) dissolved in DMSO 50% at a dose of 0.5 mg/kg/day [23, 25], using osmotic infusion pumps (Alzet Osmotic Pumps, Cupertino, CA, USA; Model 2ML4; 2.5 μL/h/28 days) implanted subcutaneously between the females’ scapulae on the gestational day 0 under anesthesia with isoflurane (Cristália, São Paulo, Brazil, 5% in 100% O_2_ for induction and 1% for maintenance) and aseptic conditions. Before and during the surgery, the anesthetic plan was checked by the absence of paw withdrawal reflex in response to a pinch stimulus and monitored by examining the animal’s respiratory frequency. Following the surgical procedure, the animals were observed until they regained consciousness. After the birth, the osmotic pumps were removed from the mothers under isoflurane anesthesia, as aforementioned. The pregnant females did not receive analgesic treatment after the procedures to implant and remove the osmotic pump to avoid potential interference with offspring development.

### 2.3 Working heart–brainstem preparation (WHBP)

Juvenile male and female offspring from the vehicle and WIN groups (P27-28) were heparinized (1,000 IU) and deeply anesthetized with isoflurane until respiratory movements ceased and withdrawal responses to noxious pinching of the tail and paw were absent. Subsequently, a transection was performed caudally to the diaphragm with the animal submerged in an ice-cold Ringer’s solution (in mM: 125 NaCl, 24 NaHCO_3_, 3.75 KCl, 2.5 CaCl_2_, 1.25 MgSO_4_, 1.25 KH_2_PO_4_, 20 glucose). The animals were then decerebrated at the precollicular level for desensitization and skinned. The lungs were removed, and the preparation was transferred to a recording chamber. The descending aorta was cannulated and retrogradely perfused using a perfusion pump (Watson-Marlow 502s, Falmouth, Cornwall, UK) through a double-lumen cannula. The perfusate consisted of Ringer’s solution containing 1.25% polyethylene glycol (an oncotic agent, Sigma, St Louis, USA) and a neuromuscular blocker (vecuronium bromide, 3-4 μg/mL, Cristália Produtos Químicos Farm. Ltda., São Paulo, Brazil). The perfusate was continuously bubbled with carbogen (5% CO_2_ and 95% O_2_, pH ∼7.4), warmed to 31–32°C (temperature measured at the point of entry into the aorta), and filtered using a nylon mesh (pore size: 25 μm, Millipore, Billirica, MA, USA). The perfusion pressure was maintained between 50–70 mmHg by adjusting the flow between 21–25 mL/min and by adding vasopressin (0.6–1.2 nM, Sigma, St. Louis, MO, EUA) to the perfusate, as previously described [26–28].

### 2.4 Nerve recordings

Multiple respiratory and sympathetic nerves were dissected, and their activities were recorded simultaneously (Fig. 1) using bipolar glass suction electrodes held in micromanipulators (Narishige, Tokyo, Japan). The inspiratory motor output was accessed through recordings of left phrenic nerve (PN) activity, with its rhythmic ramping discharge pattern used to monitor the preparation’s viability [29]. The left vagus nerve (cVN) was isolated at the cervical level and cut, and its central activity was recorded to measure the motor output to the laryngeal abductor and adductor muscles. The thoracic sympathetic chain (tSN) was dissected at T10-T12 and sectioned to record its activity. Motor output to abdominal muscles (AbN) was monitored by measuring the activity from the right lumbar plexus at the thoracic-lumbar level (T12–L1). All the signals were amplified, band-pass filtered (0.1-3 kHz, Grass Technologies, Middleton, USA), and acquired in an A/D converter (CED 1401, Cambridge Electronic Design, CED, Cambridge, UK, sampling rate: 6KHz) to a computer using Spike 2 software (version 7.1, CED).

**Figure 1.**
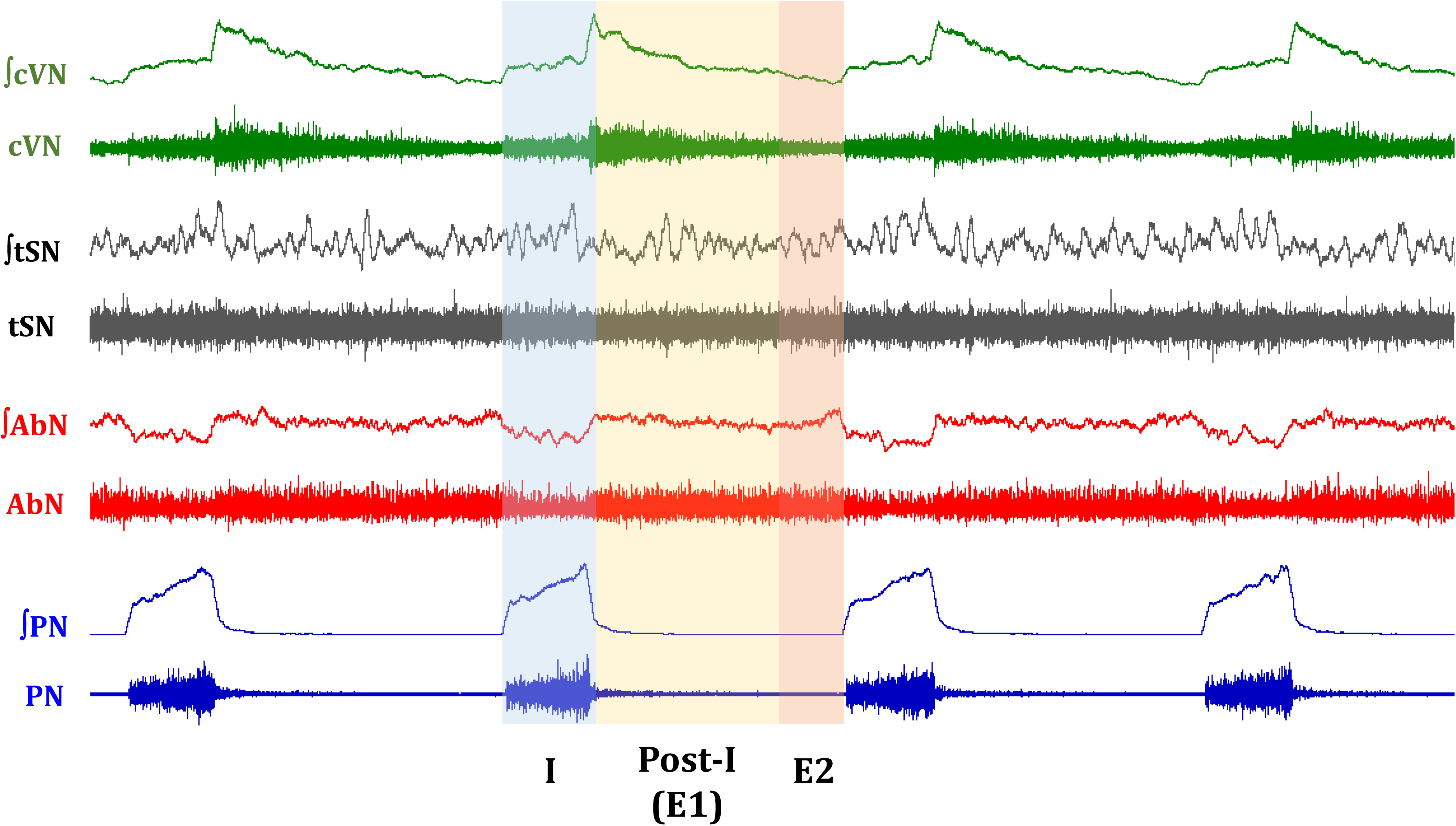
Representative raw and integrated (∫) recordings of cervical vagus nerve (cVN), thoracic sympathetic nerve (tSN), abdominal nerve (AbN), and phrenic nerve (PN) in a juvenile vehicle male illustrating the activity pattern under baseline conditions. The inspiration is highlighted by the blue area (I), passive expiration by the yellow area (Post-I / E1), and late expiration by the red area (E2).

### 2.5 Experimental protocol

After the preparation stabilization (30-40 minutes after the initiation of the perfusion), baseline PN, cVN, AbN, and tSN activities were recorded for 10-15 minutes. Then, the peripheral chemoreceptors were stimulated with potassium cyanide (KCN 0.05 %, 50 µL, dissolved in water, Sigma Chemical CO) administrated intra-arterially through a syringe connected to the perfusion system. After a 15-minute recovery period, the CO_2_ concentration in the perfusate was increased to 8% (balanced with O_2_) for 5 minutes to simulate a hypercapnic condition, followed by a 20-minute recovery period. The peripheral chemoreceptor stimulation and hypercapnia were applied randomly among experiments. At the end of the experiments, the perfusion pump was turned off to cause the preparation’s death and record the electrical noise.

### 2.6 Data analysis

Analyses were conducted offline on the rectified and smoothed signals (100 ms, Fig. 1) using custom-written scripts in Spike 2 software (Cambridge Electronic Design), as previously described [28, 30, 31]. The PN activity was analyzed by its burst amplitude (µV), frequency (time interval between consecutive bursts, expressed as cycles per minute, cpm), duration (T_I_: time of inspiration, s), and interval (T_E_: time of expiration, s). For the cVN, the activities of the inspiratory (corresponding to PN burst) and post-inspiratory (remaining activity during the expiratory phase) components were averaged and expressed as µV. The duration of the cVN post-inspiratory component was also measured and expressed either in seconds or as a percentage of T_E_. The AbN output was quantified as the mean activity (µV) during the initial two-thirds of the expiratory cycle (E1 phase, corresponding to the post-inspiratory phase) and at the final one-third (E2 phase). The tSN activity was assessed as the average total activity (µV), or the mean activity during each respiratory phase: inspiration, E1, and E2 (Fig. 1). The respiratory and sympathetic changes elicited by the peripheral chemoreceptor stimulation were quantified as the maximal changes noted during 3–5 respiratory cycles after KCN injection, corresponding to the response peak. The alterations induced by hypercapnia were evaluated using 30-second intervals during the maximal responses, observed during the final 1-2 minutes of exposure. The responses were expressed in absolute units or percentage changes relative to the baseline activities before the stimuli.

The results are reported as mean ± SD and graphically presented as boxplots, showing the median, 25%, and 75% quartiles, with individual values. GraphPad Prism version 10 software was used for graphical and statistical analyses. The data normal distribution was initially checked using the Shapiro-Wilk test. Respiratory and sympathetic baseline parameters were compared between vehicle and WIN-treated groups within the same sex using unpaired Student t-test. The responses to KCN and high CO_2_ exposure were compared between groups using either the analyses of variance for two-factor comparisons (repeated measurements two-way ANOVA), followed by Bonferroni’s post-hoc test (when baseline vs condition was considered) or unpaired Student t-test (when the magnitude of the KCN or CO_2_-induced responses was considered). Statistical results with *P* < 0.05 were considered significant.

## 3. RESULTS

### 3.1 Basal respiratory and sympathetic motor outputs

Table 1 presents the data for all nerve recordings under baseline conditions in juvenile male and female animals. No significant differences were noted in the PN activity of males treated with WIN (n=9) or vehicle (n=10), albeit the WIN-treated group showed a marginal but not statistical (*P* = 0.056) increase in the PN burst amplitude (Supplementary Fig. 1A). As for females, the WIN group (n=7) showed a shorter duration of inspiration under baseline conditions (*P* = 0.043) than the vehicle group (n=6), with no differences in the PN burst frequency, amplitude, and expiratory duration.

**Table 1.** Average values of phrenic nerve (PN) frequency, amplitude, time of inspiration (TI) and expiration (TE); cervical vagus (cVN) mean activity, peak and duration (total and relative percentage of total expiratory time - TE) at post-inspiratory phase; abdominal nerve (AbN) activity during first (E1) and second (E2) stages of expiration; and thoracic sympathetic nerve (tSN) total activity during inspiration, E1, and E2 stages of expiration for vehicle and WIN-treated juvenile male and female rats under basal conditions.

|  |  | MALE |  | P value | FEMALE |  | P value |
| --- | --- | --- | --- | --- | --- | --- | --- |
|  |  | Vehicle (n=10) | WIN (n=9) |  | Vehicle (n=6) | WIN (n=7) |  |
| PN | frequency (cpm) | 14 ± 3 | 16 ± 4 | $P = 0.328$ | 18 ± 4 | 21 ± 5 | $P = 0.259$ |
| | amplitude (μV) | 169.4 ± 43.9 | 238.4 ± 96.1 | $P = 0.056$ | 191.3 ± 121.8 | 192.9 ± 114.6 | $P = 0.982$ |
| | TI (s) | 0.79 ± 0.12 | 0.82 ± 0.26 | $P = 0.806$ | 0.93 ± 0.17 | 0.73 ± 0.14* | **$P = 0.043$** |
| | TE (s) | 3.6 ± 0.9 | 3.1 ± 0.8 | $P = 0.222$ | 2.6 ± 0.8 | 2.3 ± 0.7 | $P = 0.524$ |
|  |  | Vehicle (n=6) | WIN (n=7) |  | Vehicle (n=3) | WIN (n=4) |  |
| cVN | total activity (μV) | 23.7 ± 6.2 | 23.9 ± 8.7 | $P = 0.977$ | 35.1 ± 1.8 | 28.9 ± 1.5* | **$P = 0.004$** |
| | post-I peak (μV) | 28.5 ± 5.6 | 34.3 ± 19.9 | $P = 0.500$ | 62.9 ± 10.2 | 40.8 ± 0.4* | **$P = 0.007$** |
| | post-I duration (s) | 2.8 ± 1.2 | 2.2 ± 0.5 | $P = 0.182$ | 2.1 ± 1.1 | 2.2 ± 0.6 | $P = 0.898$ |
| | post-I duration (% T<sub>E</sub>) | 83.1 ± 7.3 | 74.3 ± 8.5 | $P = 0.071$ | 85.6 ± 5.3 | 79.9 ± 8.0 | $P = 0.336$ |
|  |  | Vehicle (n=10) | WIN (n=9) |  | Vehicle (n=4) | WIN (n=5) |  |
| AbN | E1 (μV) | 82.0 ± 50.8 | 59.2 ± 33.6 | $P = 0.272$ | 63.7 ± 34.0 | 42.9 ± 33.9 | $P = 0.392$ |
| | E2 (μV) | 66.2 ± 64.0 | 49.8 ± 41.8 | $P = 0.523$ | 44.5 ± 24.8 | 37.3 ± 35.3 | $P = 0.742$ |
|  |  | Vehicle (n=8) | WIN (n=9) |  | Vehicle (n=6) | WIN (n=6) |  |
| tSN | total activity (μV) | 286.9 ± 192.1 | 201.3 ± 112.9 | $P = 0.274$ | 210.6 ± 206.5 | 161.4 ± 151.6 | $P = 0.648$ |
| | inspiration (μV) | 308.8 ± 215.2 | 220.8 ± 124.6 | $P = 0.311$ | 224.1 ± 228.3 | 171.0 ± 166.0 | $P = 0.655$ |
| | E1 (μV) | 298.0 ± 191.5 | 225.1 ± 125.2 | $P = 0.362$ | 230.5 ± 212.8 | 176.3 ± 160.6 | $P = 0.629$ |
| | E2 (μV) | 253.9 ± 179.6 | 158.1 ± 92.0 | $P = 0.179$ | 177.1 ± 179.3 | 136.9 ± 128.6 | $P = 0.665$ |

Regarding central vagus activity, males from WIN (n=7) and vehicle (n=6) groups showed similar total activity and post-I peak amplitude and duration values. In contrast, WIN-treated female (n=4) preparations showed a depressed cVN activity (total activity and post-I peak) compared to the vehicle (n=3) preparations (*P* = 0.004 and *P* = 0.007, respectively) (Supplementary Fig. 1B). Concerning AbN activity, both male and female vehicle (n=10 and 4) and WIN (n=9 and 5) groups showed similar low-amplitude levels during baseline conditions. Additionally, both vehicle (male: n=8; female: n=6) and WIN (male: n=9; female: n=6) groups showed similar total and respiratory-modulated tSN levels, regardless of sex.

### 3.2 Respiratory and sympathetic responses to peripheral chemoreflex stimulation

Figure 2 shows PN frequency, amplitude, T_I_, and T_E_ for males (A) and females (B) during chemical activation of carotid body (CB) peripheral chemoreceptors. Peripheral chemoreceptor stimulation increased PN burst frequency (*P* < 0.0001, for both groups; Fig. 2A1) and amplitude (WIN: *P* = 0.006; Fig. 2A2) and reduced the duration of inspiratory (vehicle: *P* < 0.0001; WIN: *P =* 0.004; Fig. 2A3) and expiratory phases (*P* < 0.0001, for both groups; Fig. 2A4) in the vehicle and WIN-treated male groups. However, the shortening in T_I_ was attenuated in the WIN group compared to vehicle preparations (Δvehicle: −0.19 ± 0.03 *vs* ΔWIN: −0.10 ± 0.02 s, *P* = 0.03) (Fig. 2A3 inset and Supplementary Fig. 1C). Regarding the female groups, CB stimulation with KCN also increased PN burst frequency (*P* = 0.003, for both groups; Fig. 2B1) and amplitude (vehicle: *P* = 0.002; WIN: *P* = 0.01; Fig. 2B2), while decreasing T_E_ (*P* = 0.007, for both groups; Fig. 2B4) in vehicle and WIN-treated groups. Peripheral chemoreflex activation reduced T_I_ only in the vehicle group (*P* = 0.02), while the WIN group was unable to reduce the duration of inspiration during peripheral chemoreceptor stimulation (Δvehicle: −0.22 ± 0.06 *vs* ΔWIN: 0.02 ± 0.08 s, *P* = 0.03; Fig. 2B3).

**Figure 2.**
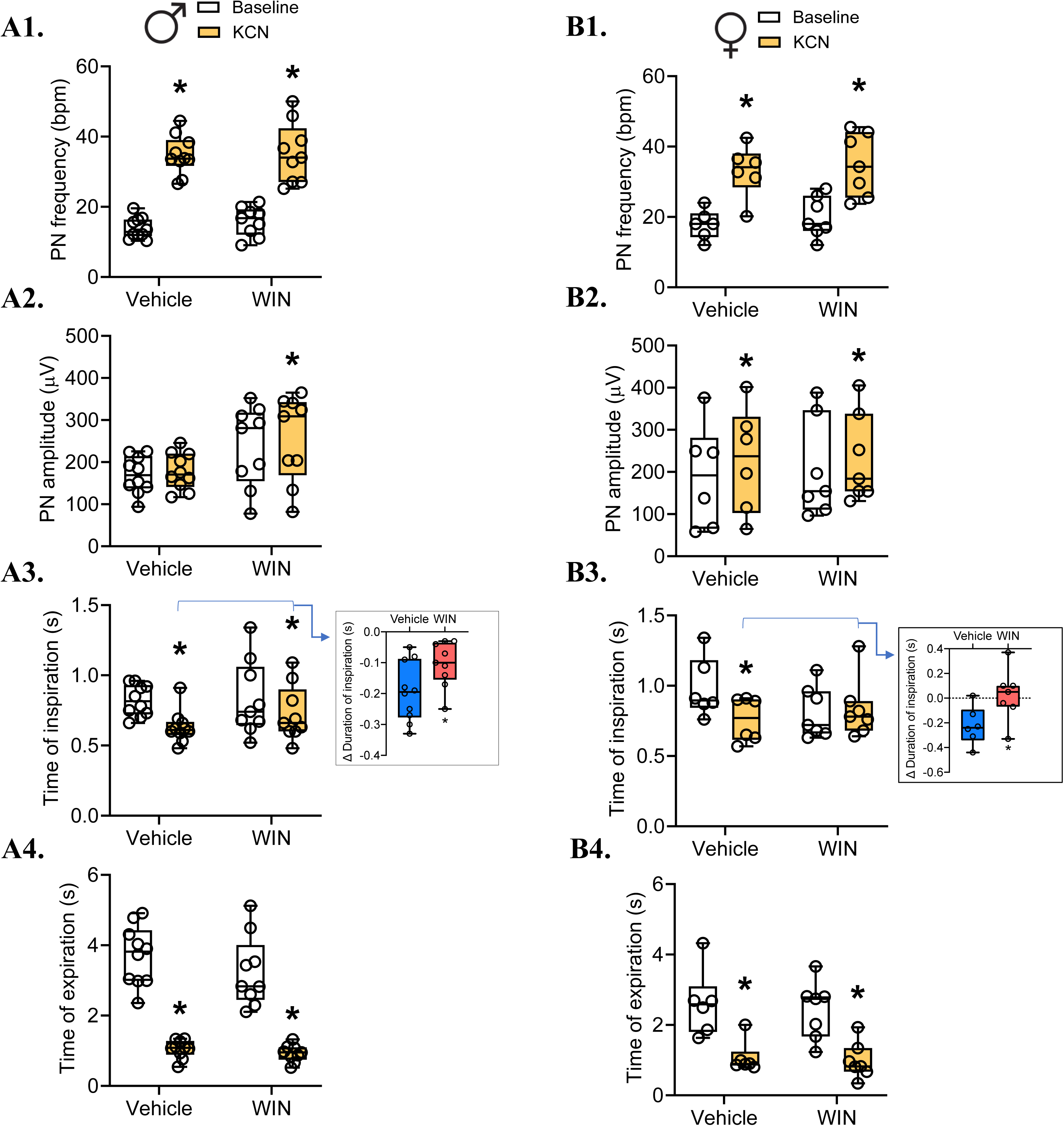
Average values of phrenic nerve (PN) frequency, amplitude, time of inspiration and expiration in juvenile male (A - Vehicle: n = 10; WIN: n = 9) and female (B - Vehicle: n = 6; WIN: n = 7) rat preparations under baseline condition and during stimulation of peripheral chemoreceptors with KCN. *Indicates a significant difference between baseline and KCN administration within the same group and sex. The inset graphs show the percentage changes relative to baseline (delta, %) in the vehicle and WIN groups.

Figure 3 illustrates the data related to cVN activity during KCN administration in males and females. In both vehicle and WIN-treated males, peripheral chemoreceptor stimulation increased the cVN discharge (*P* < 0.0001 for both groups; Fig. 3A1). KCN also elicited an increase in the post-I peak (vehicle: *P* = 0.03; WIN: *P* = 0.01; Fig. 3A2) but reduced its discharge time during expiration (*P* < 0.0001, for both groups; Fig. 3A3). The shortening of post-I duration during peripheral chemoreceptor activation was attenuated in the male WIN group (Δvehicle: −2.39 ± 0.35 *vs* ΔWIN: −1.55 ± 0.20 s, *P* < 0.05) (Fig. 3A3 inset and Supplementary Fig. 2). Regarding the females, both vehicle and WIN-treated animals increased the overall discharge of the cVN during the activation of peripheral chemoreceptors (vehicle: *P* = 0.0002; WIN: *P* = 0.003; Fig. 3B1). However, the magnitude of this response was significantly lower in the WIN group (Δvehicle: 80.0 ± 10.8 *vs* ΔWIN: 38.2 ± 7.4 %, *P* = 0.02; Fig. 3B1 inset). Likewise, the increase in the peak of cVN activity elicited by KCN administration was only noticed in the vehicle group (*P* = 0.01), with no effect in WIN-treated females (Fig. 3B2). The reduction in cVN activity within the post-I component after KCN administration was similar between female groups (Fig. 3B3).

**Figure 3.**
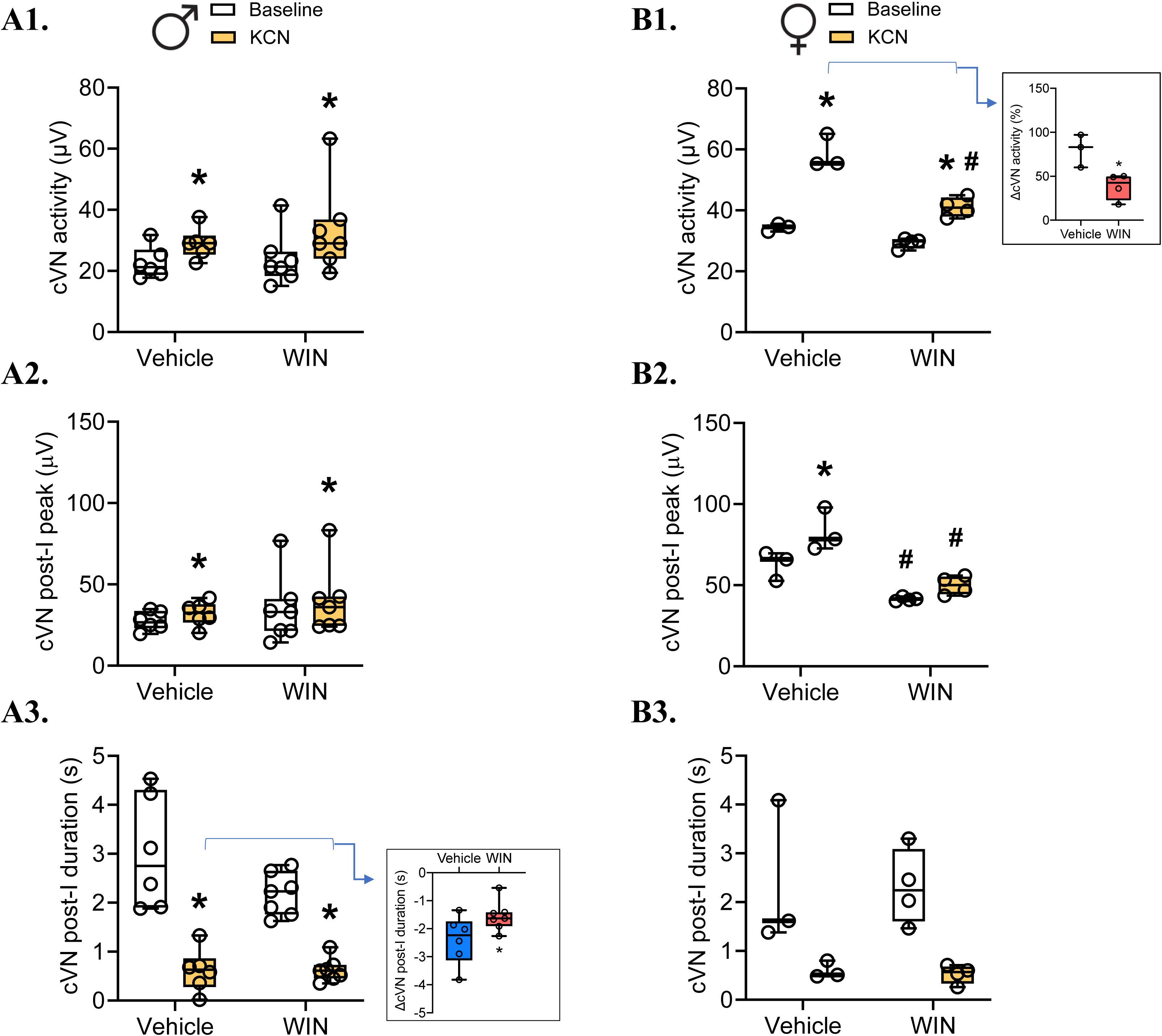
Average values of central vagus (cVN) mean activity, post-inspiratory peak and duration in juvenile male (A - Vehicle: n = 6; WIN: n = 7) and female (B - Vehicle: n = 3; WIN: n = 4) rats under baseline conditions and during stimulation of peripheral chemoreceptors with KCN. *Indicates a significant difference between baseline and KCN administration within the same group and sex. **^#^**Statistically different from vehicle animals in the same condition within the sex. The inset graphs show the percentage changes relative to baseline (delta, %) in the vehicle and WIN groups.

Figure 4 shows the data for AbN activity during the E1 and E2 stages of expiration during peripheral chemoreceptor stimulation in males (A) and females (B). KCN elicited marked increases in AbN activity in males. In the vehicle group, AbN discharge increased during E1 (*P* < 0.0001; Fig. 4A1) and E2 stages (*P* = 0.0005; Fig. 4A2). In the WIN group, the AbN response was mainly present in the E1 phase (*P* = 0.02; Fig. 4A1), with no significant increase in the activity during the E2 phase (Δvehicle: 1,080.1 ± 306.2 *vs* ΔWIN: 376.2 ± 77.6 %, *P* = 0.04; Fig. 4A2 inset). When comparing male groups (Fig. 4A), the increases in AbN activity during peripheral chemoreceptor stimulation were suppressed in the WIN group compared to the vehicle group (Supplementary Fig. 3B). Similar to the males, the AbN discharge also increased during

**Figure 4.**
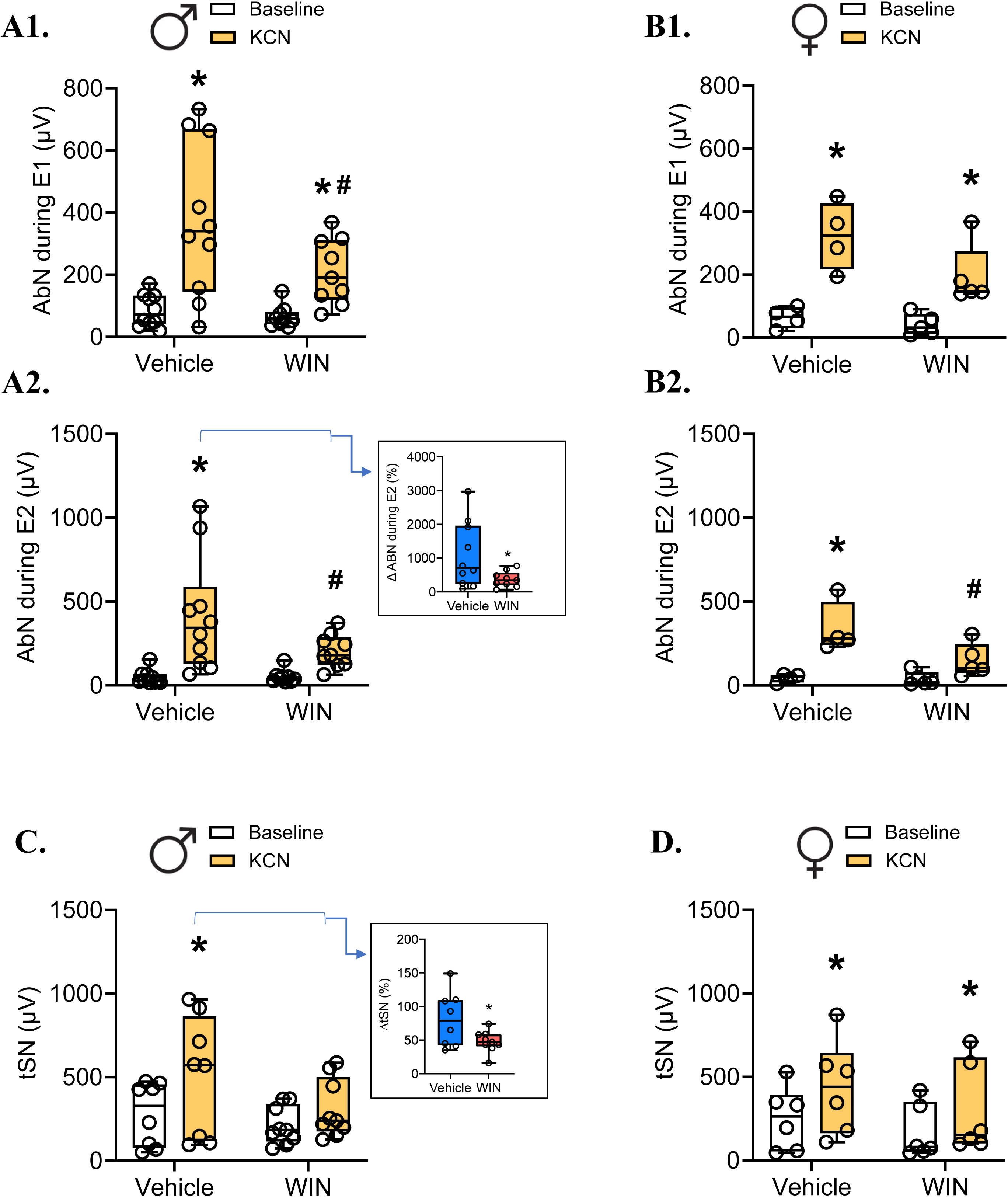
Average values of abdominal nerve (AbN) activity during first (E1) and second (E2) stages of expiration in juvenile male (A - Vehicle: n = 10; WIN: n = 8) and female (B - Vehicle: n = 4; WIN: n = 5) rat preparations, and average thoracic sympathetic nerve (tSN) activity in males (C - Vehicle: n = 8; WIN: n = 9) and females (D - Vehicle: n = 6; WIN: n = 6) under baseline conditions and during peripheral chemoreceptor stimulation with KCN. *Indicates a significant difference between baseline and KCN administration within the same group and sex. **^#^**Statistically different from vehicle animals in the same condition within the sex. The inset graphs show the percentage changes relative to baseline (delta, %) in the vehicle and WIN groups.

the activation of the peripheral chemoreceptors with KCN in the females (Fig. 4B). In the vehicle group, AbN activity increased during the E1 and E2 stages of expiration (*P* = 0.001, for both stages; Fig. 4B1 and B2, respectively). In the WIN group, the AbN response occurred mainly during the E1 phase (*P* = 0.001), with no significant changes in the activity during the E2 phase. The KCN-induced AbN activity was marginally lower during the E1 phase (*P* = 0.053) and significantly attenuated during the E2 phase (*P* = 0.01) in the WIN group.

Figure 4 also shows the results for tSN changes during peripheral chemoreceptor stimulation in males (C) and females (D). In the male groups, KCN administration evoked a significant increase in tSN activity only in the vehicle group (*P* = 0.0005; Fig. 4C). As a result, the KCN-induced sympatho-excitation in the WIN group was attenuated compared to the vehicle group (Δvehicle: 80.7 ± 14.3 *vs* ΔWIN: 47.6 ± 5.3 %, *P* = 0.04) (Fig. 4C inset and Supplementary Fig. 3C). Regarding the females, both groups displayed similar sympatho-excitatory responses to CB stimulation with KCN (vehicle: *P* = 0.003; WIN: *P* = 0.02; Fig. 4D).

### 3.3 Respiratory and sympathetic changes induced by exposure to high CO_2_ levels

Figure 5 shows the changes in PN frequency, amplitude, T_I_, and T_E_ of male (A) and female (B) groups during hypercapnic conditions (8% CO_2_). Hypercapnia exposure caused a modest but significant increase in the PN burst frequency of the vehicle group (*P* = 0.03) but not in the WIN male group (Fig. 5A1). No changes were noted in the PN burst amplitude of both groups (Fig. 5A2). Moreover, high CO_2_ exposure similarly reduced the duration of inspiration (vehicle: *P* < 0.0001; WIN: *P* = 0.002; Fig. 5A3) but not expiration (Fig. 5A4) in both male groups. In females, hypercapnia exposure did not alter PN burst frequency in both groups (Fig. 5B1). During high CO_2_ levels, PN burst amplitude increased only in the WIN group (*P* = 0.04; Fig. 5B2), while the duration of inspiration decreased only in the female vehicle group, both in absolute (*P* = 0.006) and percentage variation relative to baseline (Δvehicle: −0.17 ± 0.02 *vs* ΔWIN: 0.005 ± 0.05 s, *P* = 0.01; Fig. 5B3 and Supplementary Fig. 4A). The T_E_ was not affected by CO_2_ in either female group (Fig. 5B4).

**Figure 5.**
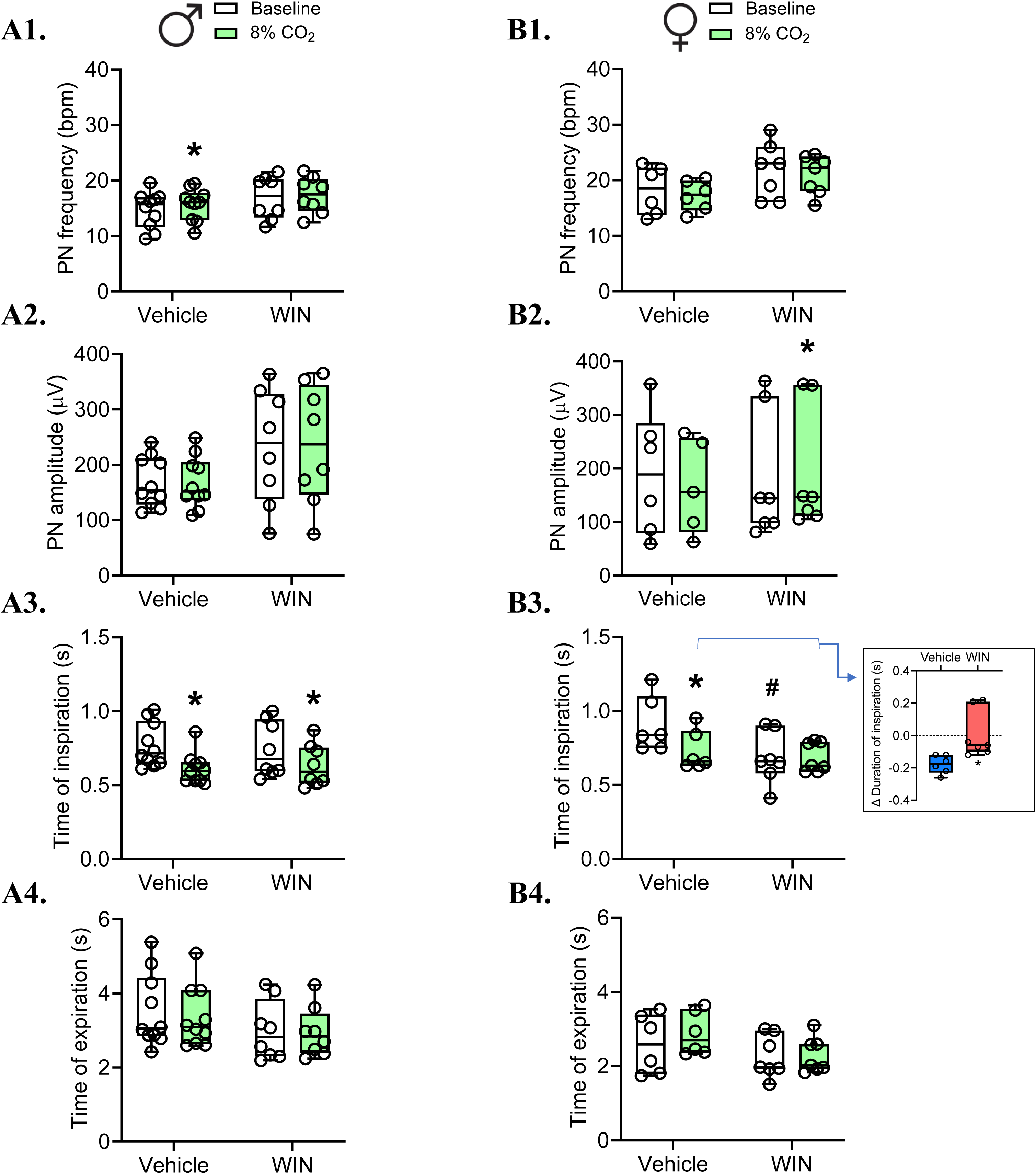
Phrenic nerve (PN) frequency, amplitude, and times of inspiration and expiration in juvenile male (A - Vehicle: n = 10; WIN: n = 8) and female (B - Vehicle: n = 6; WIN: n = 7) rats under baseline conditions and during exposure to 8% CO_2_. *Indicates a significant difference between baseline and CO_2_ exposure within the same group and sex. **^#^**Statistically different from vehicle animals in the same condition within the sex. The inset graphs show the percentage changes relative to baseline (delta, %) in the vehicle and WIN groups.

Regarding cVN activity, Figure 6 shows the data for total activity, post-I peak, and duration during high CO_2_ stimulation in males and females. In males, hypercapnia caused a modest but significant reduction in the post-I peak of both male groups (vehicle: *P* = 0.004; WIN: *P* = 0.03; Fig. 6A2), which did not affect the overall vagal activity (Fig. 6A1). High CO_2_ exposure also reduced the time of post-I discharge in the vehicle male group (*P* < 0.05) but not in the WIN group (Fig. 6A3). In females, hypercapnia increased the total mean cVN activity in the vehicle females only (*P* = 0.03), having no effects in the WIN group (Fig. 6B1 and Supplementary Fig. 4B), as also evidenced by delta analysis (Δvehicle: 25.0 ± 10.4 *vs* ΔWIN: 0 ± 3.4 %, *P* = 0.04; Fig. 6B1 inset).

**Figure 6.**
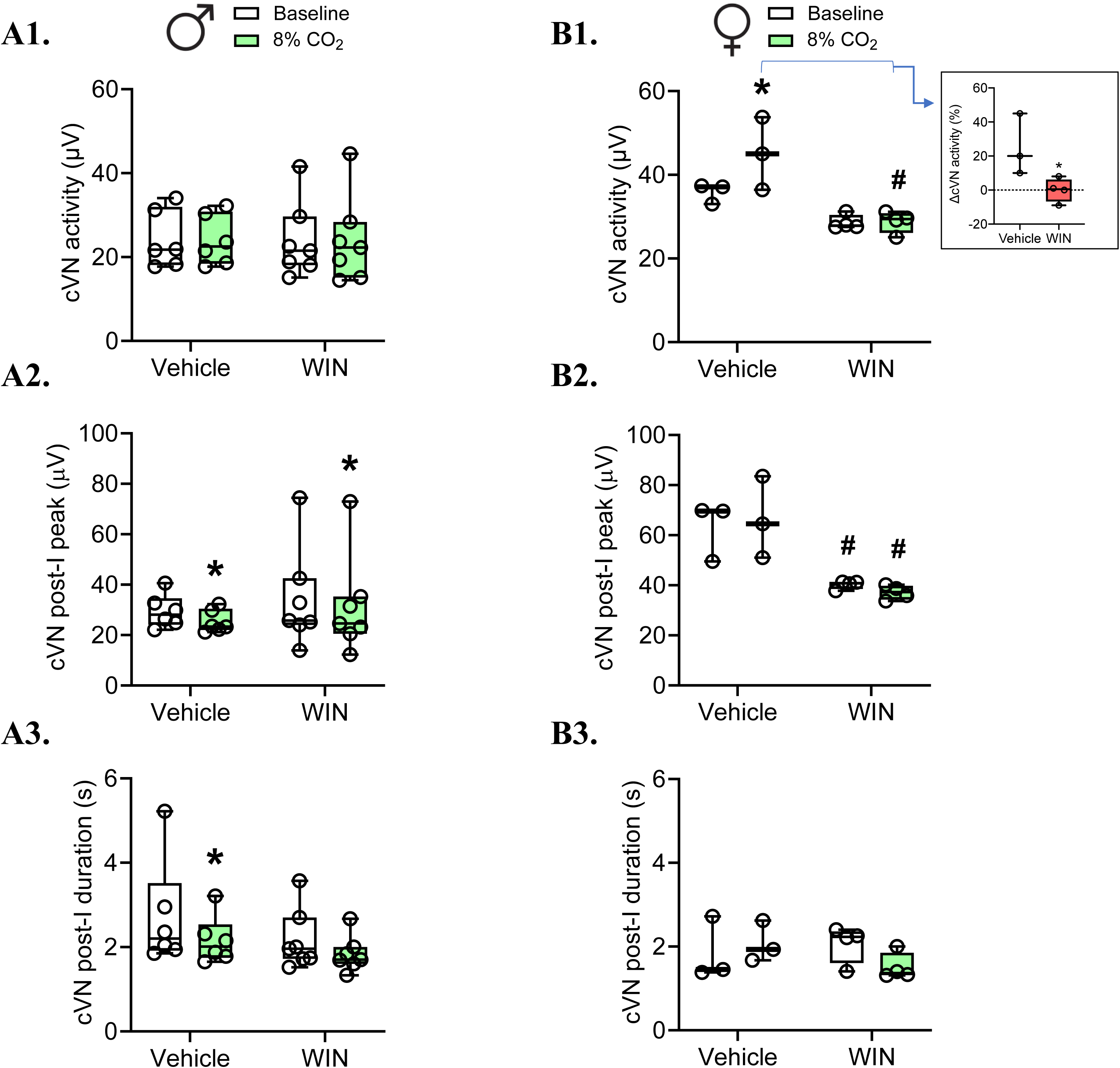
Average values of central vagus (cVN) mean activity, post-inspiratory peak and duration in juvenile male (A - Vehicle: n = 6; WIN: n = 7) and female (B - Vehicle: n = 3; WIN: n = 4) rats under baseline condition and during exposure to 8% CO_2_. *Indicates a significant difference between baseline and CO_2_ exposure within the same group and sex. **^#^**Statistically different from vehicle animals in the same condition within the sex. The inset graphs show the percentage changes relative to baseline (delta, %) in the vehicle and WIN groups.

Figure 7 represents AbN activity data during the E1 and E2 stages of expiration during hypercapnia exposure for male (A) and female (B) juvenile rats. The increase in CO_2_ concentrations brought about high amplitude bursts in AbN during the E2 phase in preparations from the vehicle male group (*P* < 0.0001; Fig. 7A2). In the WIN group, the hypercapnia-induced AbN E2 activity was marginal (*P* = 0.053) and smaller than the vehicle response (Fig. 7A2 and Supplementary Fig. 5A). As for the females, hypercapnia exposure evoked AbN E2 bursts in preparations of both groups, with similar magnitudes (vehicle: *P* = 0.006; WIN: *P* = 0.03; Fig. 7B2).

**Figure 7.**
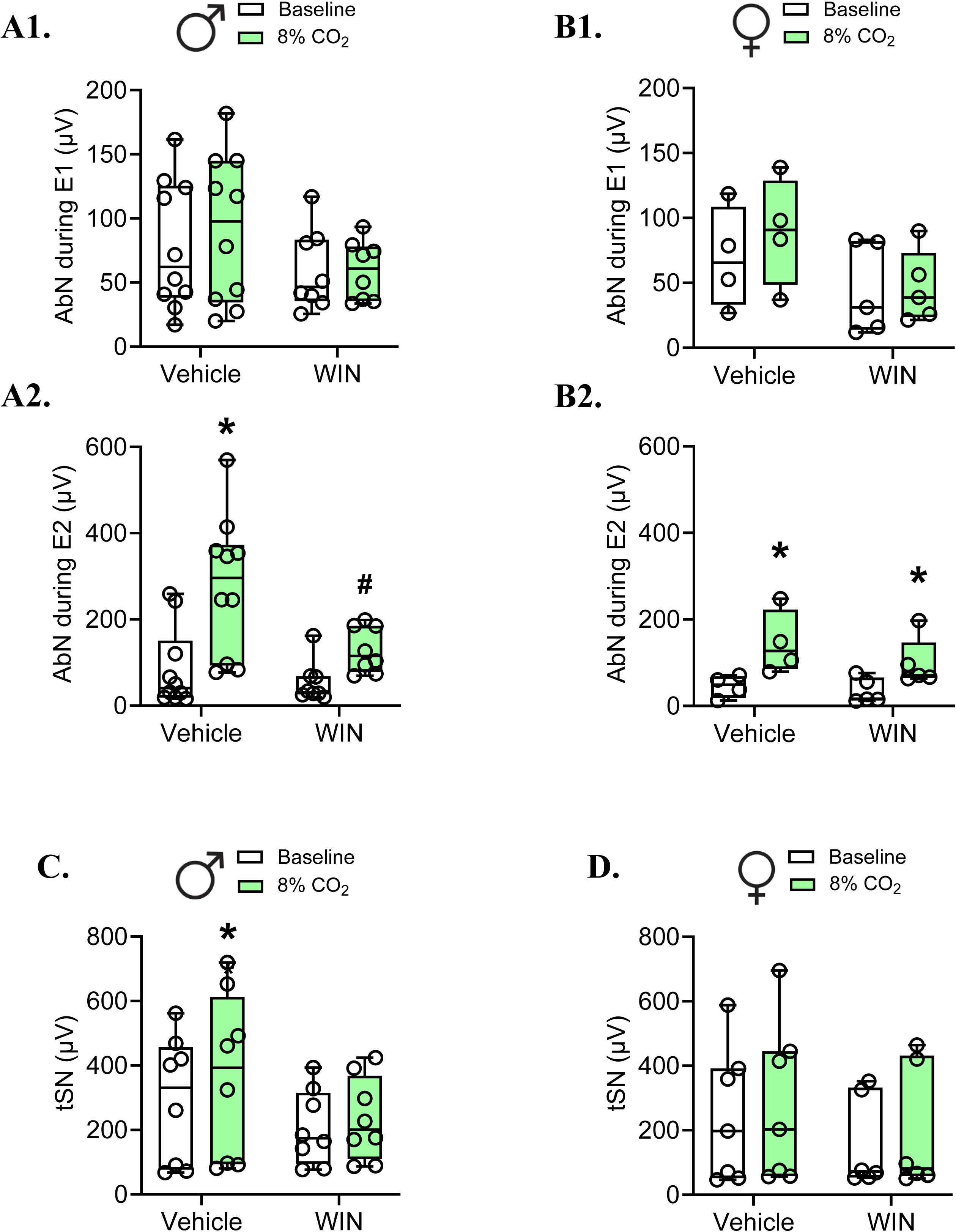
Abdominal nerve (AbN) activity during the first (E1) and second (E2) stages of expiration in juvenile male (A - Vehicle: n = 10; WIN: n = 8) and female preparations (B - Vehicle: n = 4; WIN: n = 5), and thoracic sympathetic nerve (tSN) activity in males (C - Vehicle: n = 8; WIN: n = 8) and females (D - Vehicle: n = 7; WIN: n = 6) rats under baseline condition and during exposure to 8% CO2. *Indicates a significant difference between baseline and CO_2_ exposure within the same group and sex. ^#^Statistically different from vehicle animals in the same condition within the sex.

In relation to the tSN activity, Figures 7C and D represent the data for males and females under hypercapnic conditions. CO_2_ exposure led to a significant increase in the tSN activity only in the vehicle male group (*P* = 0.01, Fig. 7C and Supplementary Fig. 5B). In contrast, hypercapnia exposure did not evoke significant increases in tSN of both female groups (Fig. 7D).

## 4. DISCUSSION

The processes governing the development of neural systems are complex and can be influenced by several factors in addition to genetic-determined mechanisms [1]. The prenatal period is a highly dynamic stage during which brain networks are highly susceptible to plasticity in response to exogenous insults, such as maternal drug use. These insults can remodel the ontogenetic development of the CNS and lead to maladaptive changes in neuronal circuitries, including the respiratory and autonomic networks, promoting long-lasting functional changes that can persist until adulthood [10, 32]. In the current study, we provide evidence showing that *in utero* exposure to synthetic cannabinoids reduces the respiratory motor drive to the upper airways and inspiratory/expiratory pumping muscles and the sympathetic discharge to blood vessels in early adult animals during blood gas disturbances. These changes elicited by prenatal exposure to WIN were qualitatively more expressive in males than females, suggesting sex-dependent effects. Our study demonstrates that endocannabinoid system overstimulation during fetal development negatively impacts networks controlling respiratory and autonomic activities and hinders the generation of appropriate cardiorespiratory adjustments during conditions of low O_2_ or excessive CO_2_ levels in the arterial blood.

### Baseline respiratory and sympathetic parameters

In males treated with WIN prenatally, we noticed a marginal trend towards increased baseline PN burst amplitude, suggesting an elevated motor activity to the diaphragm. This effect could lead to an augmented inspiratory drive and elevated resting pulmonary ventilation. This possibility agrees with our previous *in vivo* study demonstrating that juvenile male rats exposed to prenatal WIN showed baseline hyperventilation due to a higher tidal volume [23]. The increased motor drive to the diaphragm could also contribute to compensating for reduced lung compliance observed in juvenile male rats prenatally treated with WIN [23], preventing long-term alterations in pulmonary mechanisms and maintaining pulmonary ventilation in adult life [24]. Unlike males, the female group treated with WIN during the gestational period displayed reduced inspiratory phase (TI) duration under baseline conditions, with no PN burst amplitude and frequency changes. In our previous studies, we did not identify relevant alterations in resting breathing and lung compliance of juvenile [23] and adult [24] females exposed to WIN prenatally. Therefore, the mechanical impact of shorter inspiratory duration on the pulmonary ventilation of WIN-treated females still requires additional studies to be fully understood. However, these observations indicate that the control of inspiratory motor activity is also altered in this group. Endocannabinoid receptors have been identified in the ventral medulla [33] – a region containing neurons essential for respiratory rhythm and pattern generation [30, 45]. Moreover, systemic administration of the endocannabinoid anandamide in brainstem-spinal cord *en bloc* preparations of newborn mice depressed resting PN activity [34]. Based on these observations and given that the peripheral afferents influencing inspiratory motor activity, such as lung vagal afferents, are absent in *in situ* preparations, we speculate that the increased phrenic activity of phrenic in male animals and reduced inspiratory phase duration in females treated with WIN is due to persistent changes in the activity of premotor and motoneurons located in the brainstem or spinal cord.

Intrauterine exposure to the synthetic cannabinoid reduced vagal efferent activity under basal conditions in preparations of juvenile females. The vagus nerve contains parasympathetic fibers that innervate the heart and play a role in cardiovascular control [35, 36]. However, in our previous *in vivo* studies, we did not observe changes in resting heart rate of animals treated with WIN prenatally [23, 24], suggesting that the depressed vagal output noticed in the current study may not affect the cardiac activity. The vagus nerve also presents motor fibers innervating the laryngeal muscles and controlling upper airway resistance [37]. Fibers firing during post-inspiration cause larynx adduction during the expiratory phase, hence regulating the expiratory flow and lung emptying [37]. The reduced post-inspiratory drive seen in the vagal activity of female rats prenatally treated with WIN indicates a reduced motor drive to laryngeal muscles, which may generate instabilities in the upper airways that lead or predispose to breathing irregularities or obstructive events – possibilities that require further investigation to be verified.

Although studies have reported an acute excitatory effect of cannabinoid receptor activation on sympathetic nerve discharge, mainly modulating the activity of sympathetic pre-ganglionic neurons in the ventral medulla [33, 38–40], our experiments *in situ* showed that the prenatal exposure to WIN did not promote long-lasting effects on baseline vasoconstrictor sympathetic discharge in juvenile rats of either sex. These findings also parallel *in vivo* observations demonstrating that juvenile and adult animals treated with WIN prenatally exhibit similar blood pressure levels compared to non-exposed animals [23, 24]. Therefore, the sympathetic network in the brainstem of WIN-treated animals may not exhibit long-term changes that affect baseline cardiovascular control.

### Respiratory and sympathetic responses to blood gas disturbances

During conditions of low oxygen levels in the arterial blood, the carotid body chemoreceptors are activated and trigger vigorous increases in inspiratory, expiratory, and sympathetic outputs [41]. In the current study, this condition was mimicked *in situ* through injections of potassium cyanide, which also promotes the depolarization of peripheral chemoreceptor cells in the carotid bodies. In the vehicle group, KCN injections increased the respiratory frequency in association with reductions in the duration of the inspiratory and expiratory phases. This shortening in T_I_ was attenuated in the WIN-treated male and female groups. Although this effect has not impacted the development of the tachypneic response to cyanide, these observations indicate that the mechanisms promoting the inspiratory-expiratory phase transition during hypoxic situations were affected by the treatment with WIN during the prenatal period. In addition, we observed that WIN-treated males showed larger PN burst amplitude while the females exposed to WIN prenatally displayed a suppressed vagal post-I activity during carotid body stimulation with KCN, which could be a consequence of altered baseline activities observed in these groups.

We also noticed that the increase in abdominal expiratory activity during carotid body stimulation was suppressed in males and females treated with WIN, especially during the second phase of expiration. This finding suggests that the motor activity driving forced exhalation in WIN-exposed groups is attenuated, which may negatively affect expiratory flow control and the hypoxic ventilatory response (HVR). However, our previous studies performed *in vivo* indicated that the HVR is preserved in WIN-treated juvenile animals [23]. In our *in vivo* studies, the animals were exposed to a low O_2_ environment for 20 minutes, which also caused metabolic changes that impacted pulmonary ventilation [23]. In our current *in situ* study, the KCN administration caused a rapid and brief activation of carotid body chemoreceptors. Therefore, the divergent *in vivo* and *in situ* findings indicate that juvenile animals treated with WIN prenatally struggle to cope with hypoxic situations during the initial moments of exposure, but they can compensate for this motor deficit over time with a metabolic response. Interestingly, sex-dependent alterations in the O_2_ chemoreflex were observed in WIN-treated adult animals, where females exhibited a reduced HVR while males displayed an opposite response [24] – effects that might have resulted from a combination of changes in inspiratory and expiratory motor activities observed in the juvenile age.

The sympathetic-excitatory response to carotid body stimulation was reduced in males but not females treated with WIN prenatally. This attenuation may imply an impaired capacity of WIN-treated male animals to promote proper peripheral blood flow redistribution during hypoxic situations, at least during the initial moments of exposure, affecting the maintenance of homeostasis, especially in tissues that lack energy reserves, such as the brain. This sex-related effect, combined with the alterations in the respiratory responses, suggests that males exposed to cannabinoids during gestation are more susceptible to exhibiting adverse effects on the ability to mount proper responses to hypoxia than females. The depressed reflex responses to hypoxia could result from actions of the cannabinoid on the development of peripheral (carotid bodies) and central pathways (brainstem and spinal cord) that express CB_1_ receptors [42, 43].

Similar to hypoxia, exposure to high levels of CO_2_ also evokes reflex respiratory and sympathetic responses due to the activation of chemosensitive cells in the brain [45]. In our previous *in vivo* study, we observed amplified ventilatory responses to high CO_2_ exposure in WIN-treated male and female juvenile rats [23], dependent on a larger increase in tidal volume than respiratory frequency. In the long term, adult males prenatally exposed to WIN still exhibited increased hypercapnic ventilatory responses (HcVR), whereas females showed a reduced CO_2_ response [24], reinforcing a plasticity occurrence in the ventilatory control system throughout postnatal life as a consequence of fetal cannabinoid exposure. In our current *in situ* experiments with juvenile rats, we observed minor changes in the respiratory and sympathetic responses to hypercapnia in WIN-treated animals, including attenuated abdominal and sympathetic responses in males and a modest amplification of PN burst amplitude increase, blunted shortening of inspiration, depressed vagal excitation in females. The lack of peripheral feedback information, especially from the pulmonary stretch receptors, significantly affects the HcVR of vagotomized preparations [44], such as the *in situ* preparations. Despite this limitation, the *in situ* preparations are helpful for understanding the motor pattern alterations underlying the HcVR. In control animals, the increase in PN activity, generation of forced expiration (augmented AbN activity), and reduction in vagal post-I activity (leading to reduced upper airway resistance during expiration) is suggested to contribute to increasing pulmonary ventilation under hypercapnia [45]. Therefore, the fact that we could not identify a direct link between the current *in situ* study and our previous *in vivo* findings [23] indicates that factors that are absent *in situ* might contribute to the augmented HcVR observed in unanesthetized juvenile rats treated with WIN prenatally. This observation opens the possibility to explore the effects of CB_1_ receptor activation during pregnancy on the development of other mechanisms that influence the HcVR, such as pulmonary afferents, mid- and forebrain inputs to the respiratory network, or changes in the metabolism.

Our results showing that the motor and sympathetic responses to carotid body stimulation (similar to hypoxic condition) or exposure to hypercapnia in *in situ* preparations indicate that the brainstem mechanisms processing the O_2_/CO_2_ sensory information are modified by prenatal cannabinoid exposure. Although modest, the differences observed between males and females indicate sex-related mechanisms inducing the long-term effects. We hypothesize that after sexual maturation, the elevation in plasma sex hormone levels can potentiate the differences in respiratory and sympathetic control in males and females treated with WIN prenatally, leading to more pronounced sex-related differences, as documented *in vivo* [24].

In summary, our study provides evidence regarding the deleterious effects of cannabinoid use during pregnancy on the cardiorespiratory motor nerve outputs of juvenile offspring, modifying the reflex responses to low O_2_ (mimicked by chemical carotid body stimulation) or high CO_2_ levels. The use of cannabis has increased worldwide in an indiscriminate manner, and pregnant women are becoming a significant percentage of users due to liberal government policies combined with the lack of scientific knowledge about the postnatal consequences on offspring health. Thus, this study strengthens the precautionary note that must be considered for the medicinal or recreational use of cannabis during pregnancy and advances our understanding of the postnatal life implications.

## 5. ACKNOWLEDGMENTS

We thank Euclides Roberto Secato for the excellent technical assistance.

## 8 AUTHOR CONTRIBUTIONS

**Luis Gustavo A. Patrone:** conceived and designed research, performed experiments, analyzed data, interpreted results of experiments, prepared figures, drafted manuscript, edited and revised manuscript, approved final version of manuscript. **Marlusa Karlen-Amarante:** performed experiments, analyzed data, interpreted results of experiments, edited and revised manuscript, approved final version of manuscript. **Luciane H. Gargaglioni:** conceived and designed research, drafted manuscript, edited and revised manuscript, approved final version of manuscript. **Daniel B. Zoccal:** conceived and designed research, performed experiments, analyzed data, interpreted results of experiments, prepared figures, drafted manuscript, edited and revised manuscript, approved final version of manuscript.

---

**Suppl. Figure 1.**
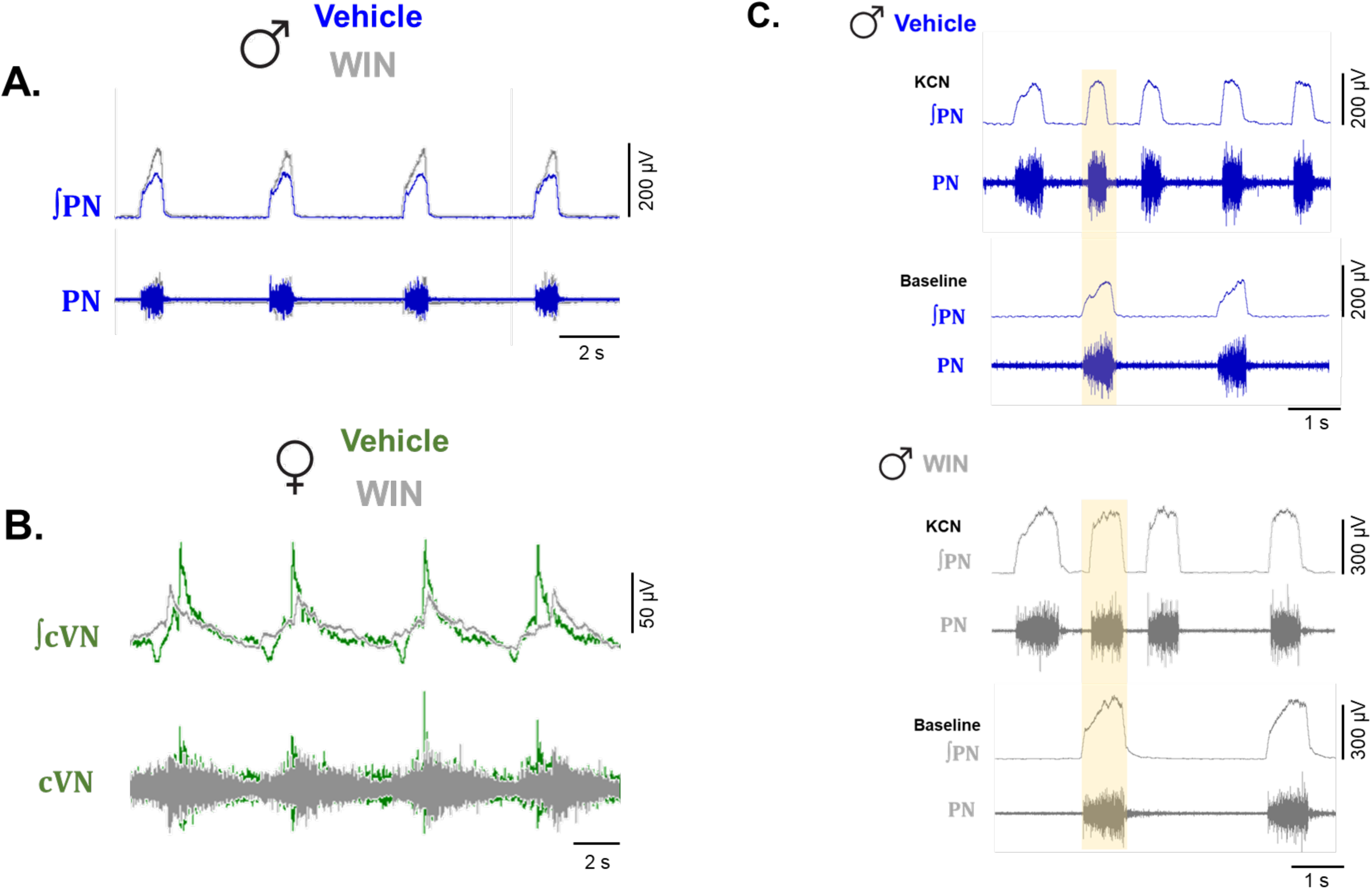
**Panel A:** Raw and integrated (∫) recordings of baseline phrenic nerve (PN) activity of representative male animals from the vehicle and WIN groups, showing an increased amplitude in treated males. **Panel B:** Raw and integrated (∫) recordings of baseline cervical vagus nerve (cVN) activity of representative females from the vehicle and WIN groups, demonstrating the reduced post-inspiratory peak activity and duration in treated females. **Panel C:** PN activity during peripheral chemoreceptor stimulation with KCN in representative vehicle and WIN males, evidencing a smaller reduction in T_I_ during KCN in the WIN group. The highlighted area corresponds to the duration of the inspiratory phase during baseline conditions.

**Suppl. Figure 2.**
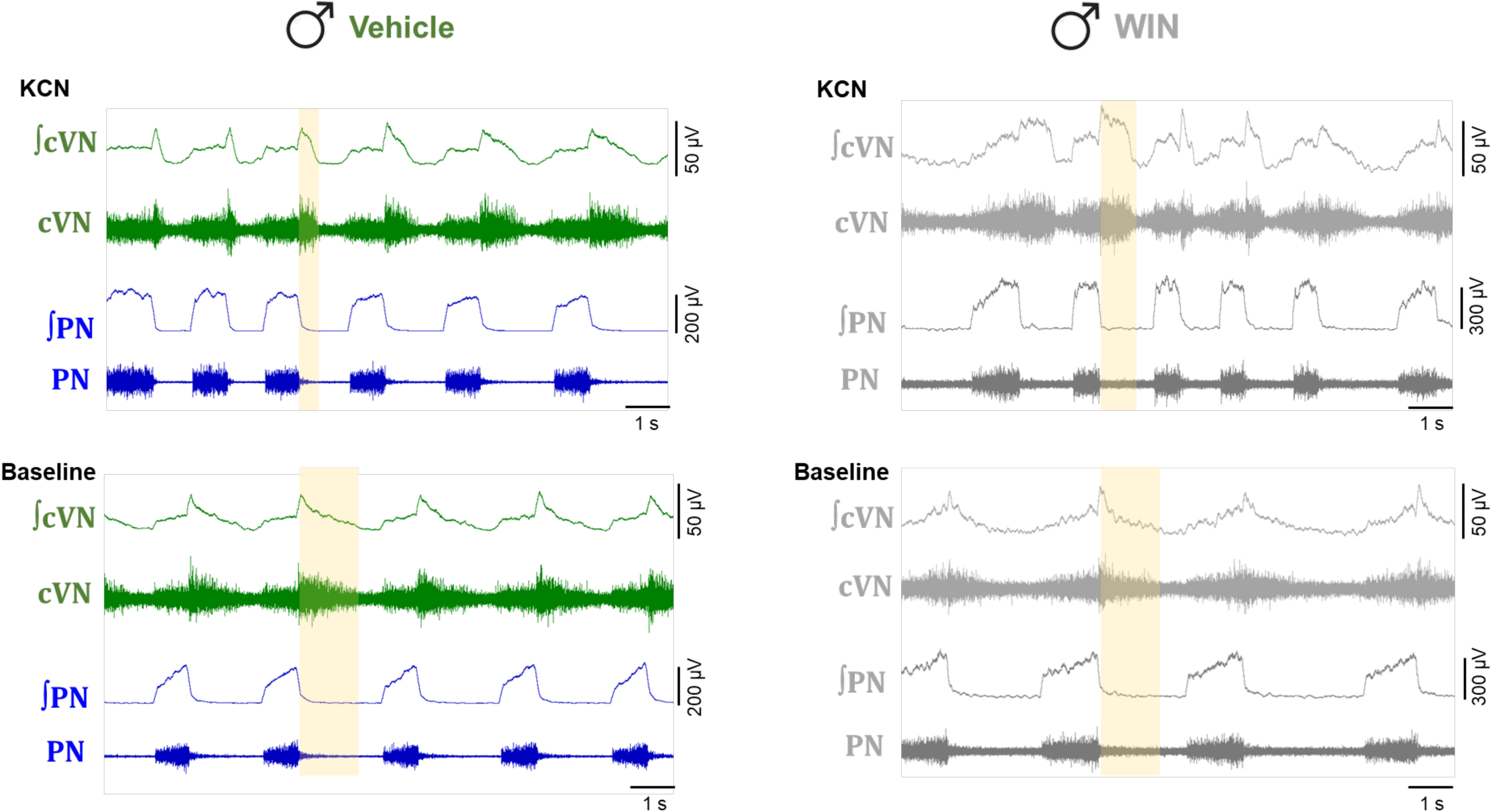
Raw and integrated (∫) recordings of cervical vagus (cVN) and phrenic nerve (PN) activities under baseline conditions and during peripheral chemoreceptor stimulation with KCN from representative vehicle and WIN males, demonstrating a smaller reduction in post-I cVN time of activity during KCN in WIN group. The highlighted area corresponds to the post-inspiration phase (E1).

**Suppl. Figure 3.**
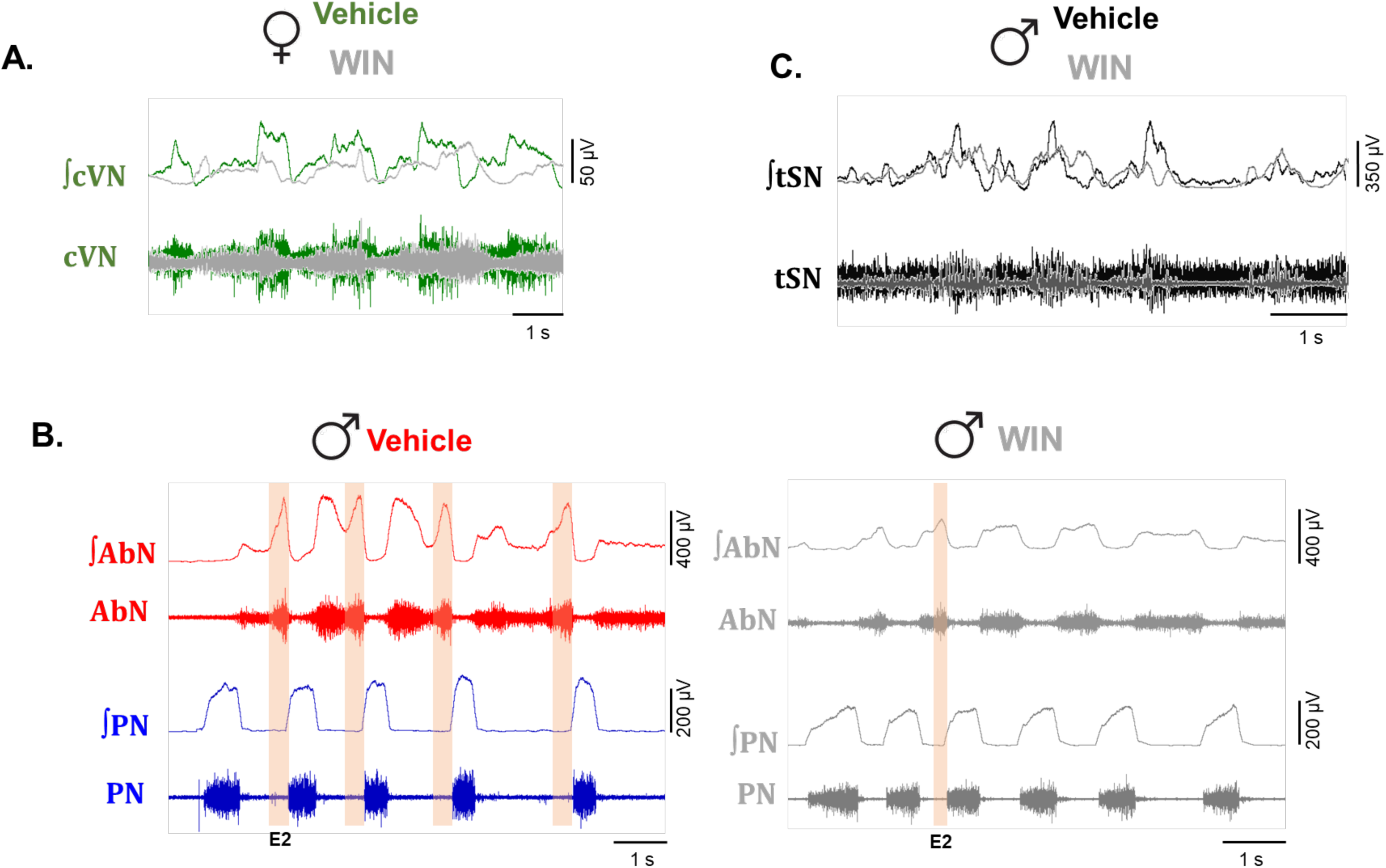
**Panel A:** Raw and integrated (∫) recordings of cervical vagus nerve (cVN) activity during KCN exposure in representative female animals from the vehicle and WIN groups, evidencing a reduced cVN response in WIN females. **Panel B:** Raw and integrated (∫) abdominal (AbN) and phrenic nerve (PN) activities during peripheral chemoreceptor stimulation with KCN in vehicle and WIN-treated male animals, demonstrating the attenuated AbN activity in WIN males. The highlighted area corresponds to the E2 phase. **Panel C:** Raw and integrated (∫) thoracic sympathetic nerve (tSN) activity during KCN exposure in representative vehicle and WIN-treated males, illustrating the lower sympatho-excitatory response in treated males.

**Suppl. Figure 4.**
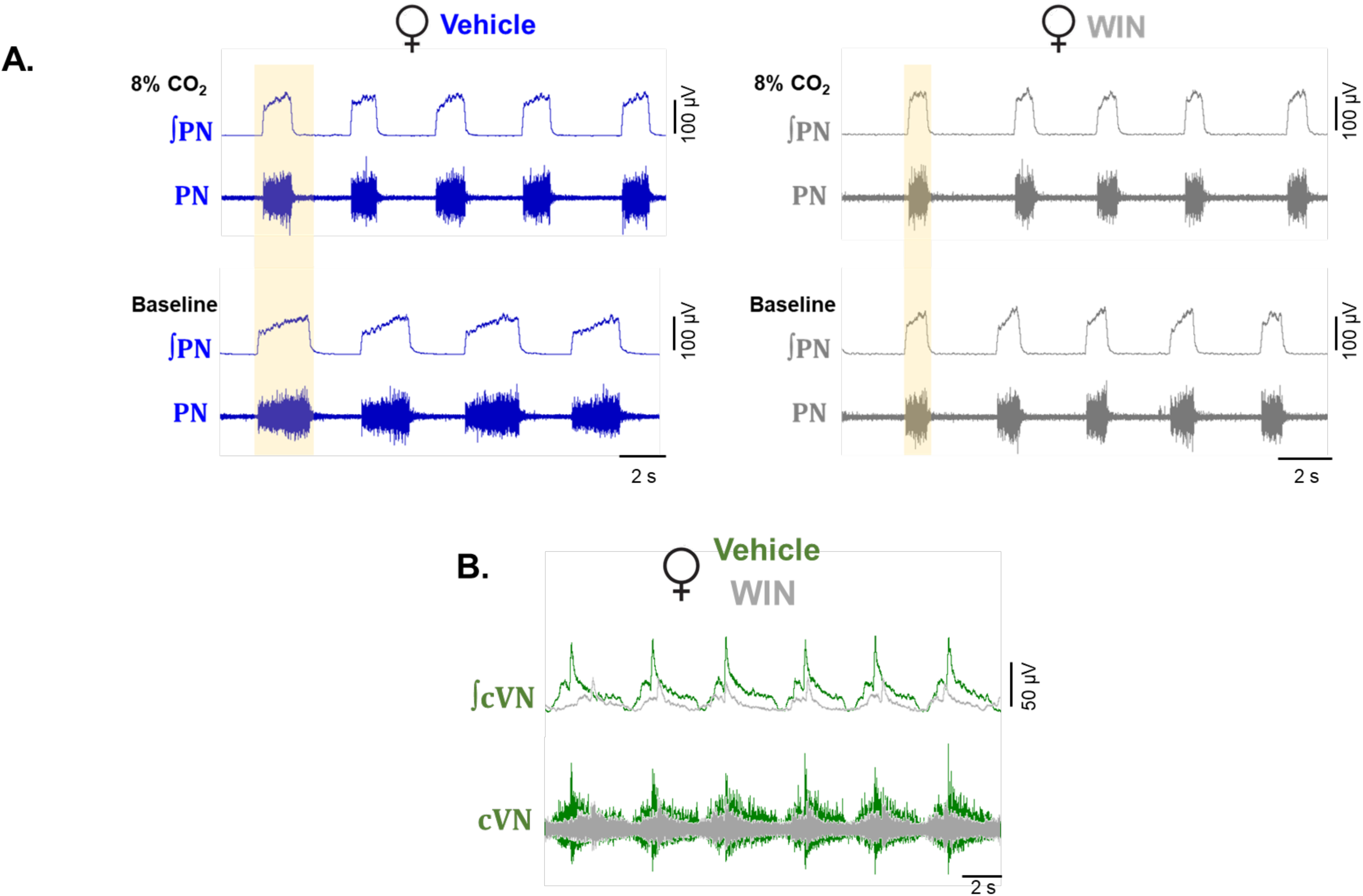
**Panel A:** Raw and integrated (∫) recordings of phrenic nerve (PN) activity under baseline conditions and during exposure to high levels of CO_2_ in representative female animals from the vehicle and WIN groups, demonstrating the lower reduction in T_I_ in treated animals during hypercapnia. The highlighted area corresponds to the duration of the inspiratory phase under baseline conditions. **Panel B:** Raw and integrated (∫) recordings of cervical vagus nerve (cVN) activity under 8% CO_2_ condition in representative vehicle and WIN females, illustrating the lower activity peaks in treated females.

**Suppl. Figure 5.**
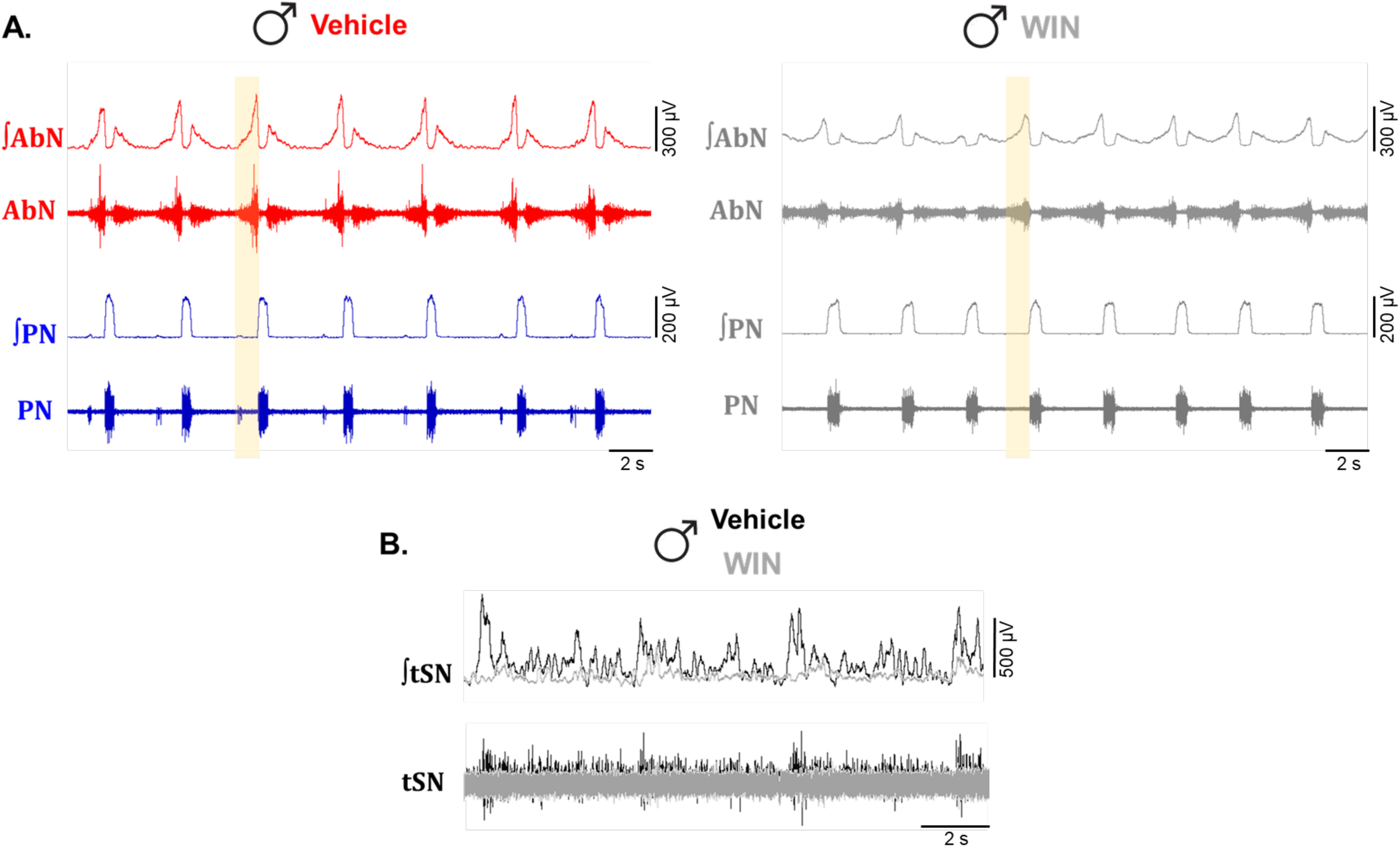
**Panel A:** Raw and integrated (∫) recordings of abdominal (AbN) and phrenic nerve (PN) activities during 8% CO_2_ exposure in representative vehicle and WIN-treated male animals, showing the lower amplitude of AbN during E2 in WIN animals. The highlighted area indicates the E2 phase. **Panel B:** Raw and integrated (∫) recordings of thoracic sympathetic nerve (tSN) activity during CO_2_ exposure in representative vehicle and WIN-treated males, demonstrating the blunted sympatho-excitatory response in the treated animals.

